# Human 28s rRNA 5’ terminal derived small RNA inhibits ribosomal protein mRNA levels

**DOI:** 10.1101/618520

**Authors:** Shuai Li

## Abstract

Recent small RNA (sRNA) high-throughput sequencing studies reveal ribosomal RNAs (rRNAs) as major resources of sRNA. By reanalyzing sRNA sequencing datasets from Gene Expression Omnibus (GEO), we identify 28s rRNA 5’ terminal derived sRNA (named 28s5-rtsRNA) as the most abundant rRNA-derived sRNAs. These 28s5-rtsRNAs show a length dynamics with identical 5’ end and different 3’ end. Through exploring sRNA sequencing datasets of different human tissues, 28s5-rtsRNA is found to be highly expressed in bladder, macrophage and skin. We also show 28s5-rtsRNA is independent of microRNA biogenesis pathway and not associated with Argonaut proteins. Overexpression of 28s5-rtsRNA could alter the 28s/18s rRNA ratio and decrease multiple ribosomal protein mRNA levels. Our results reveal that 28s5-rtsRNA serves as a key regulator in ribosomal protein expression.

## Introduction

Small RNAs (sRNAs) play diverse regulatory roles in eukaryotic cells (1–5). Recently, with the development of sRNA high-throughput sequencing technology, a big amount of novel sRNAs are identified and the category of sRNA is greatly expanded (6–11). Different housekeeping RNAs, such as snoRNA, tRNA and rRNA, could serve as sources of sRNA (12). Ample studies show that these abundant non-coding RNA-derived sRNAs play important roles. For example, sperm transfer RNA-derived sRNAs (tsRNAs) could modify offspring metabolism (13–15).

rRNA-derived sRNAs are identified by multiple studies (16). Lee *et al* found that DNA damage induced a new class of ribosomal DNA locus originated sRNAs (named qiRNA) in the filamentous fungus *Neurospora crassa* (17). In mammalian cells, rRNA-derived sRNAs exhibit special tissue distribution and are associated with inflammatory and metabolic disorder conditions (18–20). Moreover, rRNA-derived sRNAs are showed preferentially originated from the terminals of mature rRNAs (12,18).

In present research, we focus on the most abundant 5’ 28s rRNA-terminal-derived small RNA (named 28s5-rtsRNA). By analyzing published sRNA sequencing data, we find that rtsRNAs are enriched in sperm and skin tissues, knockout of Dicer or Dgcr8 (two key genes in miRNA biogenesis) could not block 28s5-rtsRNA expression. Argonaut protein RIP-sequencing (RNA-binding protein immunoprecipitation sequencing) data shows that 28s5-rtsRNA is not enriched in RISC complex. Finally, we show that overexpression of 28s5-rtsRNA significantly reduce the mRNA levels of a number of ribosomal proteins. Our results demonstrate the expression signature and the regulatory function of 28s5-rtsRNAs.

## Methods and Materials

### Cell culture

The HeLa cell line was maintained in Dulbecco’s modified Eagle’s medium (DMEM) (Thermo Fisher Scientific, Hudson, NH, USA) containing 10% fetal bovine serum (FBS) with 1% penicillin-streptomycin solution at 37°C with 5% CO_2_.

#### Small RNA high throughput sequencing data analysis

Processed sRNA sequencing datasets (contains Sequence and Count) of mouse embryonic stem cells (GSM314552) and human normal liver tissue (GSM531974) were downloaded from GEO. rRNA-derived sRNAs were identified by in-house Perl scripts. For sRNA sequencing data from Sequence Read Archive (SRA), sRNAbench tool was employed to perform data processing using default parameter (21).

### 28s5-rtsRNA mimics transfection

28s5-rtsRNA mimics (28s5-rtsRNA, sense: CGC GAC CUC AGA UCA GAC GU; antisense: GUC UGA UCU GAG GUC GGG AU) and scramble Negative Control RNA (Ctrl, sense: UUC UCC GAA CGU GUC ACG UTT; antisense: ACG UGA CAC GUU CGG AGA ATT) were synthesized by GenePharma (Shanghai, China). RNAs are transfected using Lipofectamine 2000 reagent following the protocol. Total RNA was extracted 48 h after transfection.

### RNA-sequencing and data processing

Total RNA concentration was measured using Qubit RNA Assay Kit in Qubit 2.0 Flurometer (Life Technologies, CA, USA). RNA integrity and 28s/18s ratio were assessed using the RNA Nano 6000 Assay Kit of the Bioanalyzer 2100 system (Agilent Technologies, CA, USA). Sequencing libraries were generated using NEBNext Ultra RNA Library Prep Kit for Illumina (NEB, USA) following protocol and index codes were added to attribute sequences to each sample. The library preparations were sequenced on an Illumina Hiseq platform and 150bp paired-end reads were generated. Clean data (clean reads) were obtained by removing reads containing adapter, reads containing poly-N and low quality reads from raw data. Clean reads were mapped to the reference genome using HISAT2 (v2.0.5) to generate BAM files. BAM files were visualized and analyzed using SeqMonk (v1.41.0, Babraham Bioinformatics). Gene ontology (GO) analyses were completed using ToppGene (22).

### Real-time qPCR

For mRNA expression analysis, cDNA was generated using the PrimeScript™ RT reagent Kit with gDNA Eraser (TaKaRa). cDNA was then diluted 50 times by H_2_O. Real-time PCR was performed using SYBR Premix Ex Taq™ II (Tli RNaseH Plus, TaKaRa). GAPDH mRNA levels were used for normalization. Forward (F) and reverse (R) primers used were as follows: RPS9-F 5’- ctctttctcagtgaccgggt −3’, RPS9-R 5’- tttgttccggagcccatact −3’; RPS13-F 5’- ctcctttcgttgcctgatcg −3’, RPS13-R 5’- tctgtgaaggagtaaggccc −3’; RPS14-F 5’- cgggccacaggaggaaatag −3’, RPS14-R 5’- ggtgacatcctcaatccgcc −3’; RPL14-F 5’- agaaggttcctgcccagaaa −3’, RPL14-R 5’- atctgcctcctaactccagc −3’; GAPDH-F 5’- tcagtggtggacctgacctg −3’, GAPDH-R 5’- tgctgtagccaaattcgttg −3’.

#### Statistics

For RNA-sequencing data, DESeq2 and edgeR were used to identify differential expressed genes. Realtime-qPCR relative mRNA expression levels were presented as the mean ± SD. Differences were assessed by two-tailed Student’s t-test using GraphPad software. *p* < 0.05 was considered to be statistically significant.

## Results

### rRNA derived small RNA

Previously, Li *et al* showed that housekeeping non-coding RNAs such as tRNA, snoRNA, snRNA and rRNA preferentially produce small 5’ and 3’ end fragments (12). Here we obtain two small RNA sequencing datasets from GEO (Gene Expression Omnibus), including GSM314552 of mouse embryonic stem cells (ESCs) and GSM531974 of human normal liver tissue (18,23). rRNA-derived sRNAs account for 8.13% (609589 / 7501068) and 5.65% (496998 / 8790059) of total reads in mouse ECSs and in human liver, respectively. sRNA deep sequencing reads are then aligned along with 45s pre-rRNA. As showed by Figure 1AC, 28s rRNA 5’ terminal derived small RNAs are the most abundant rRNA-derived small RNAs (accounting for 11.2% (68188 / 609589) and 17.5% (87080 / 496998) of total rRNA-derived sRNA in ESCs and in liver tissue, respectively). These 28s rRNA 5’ terminal derived small RNAs show a length dynamics with identical 5’ end and different 3’ end (Figure 1BD). In this study, we assign these sRNA into one category and name them as 28s5-rtsRNA. In mouse ESCs, the most abundant 28s5-rtsRNA is 17nt in length. While for human liver tissue, most 28s5-rtsRNAs are enriched within the range of 19-21nt (Figure 1BD).

**Figure 1.**
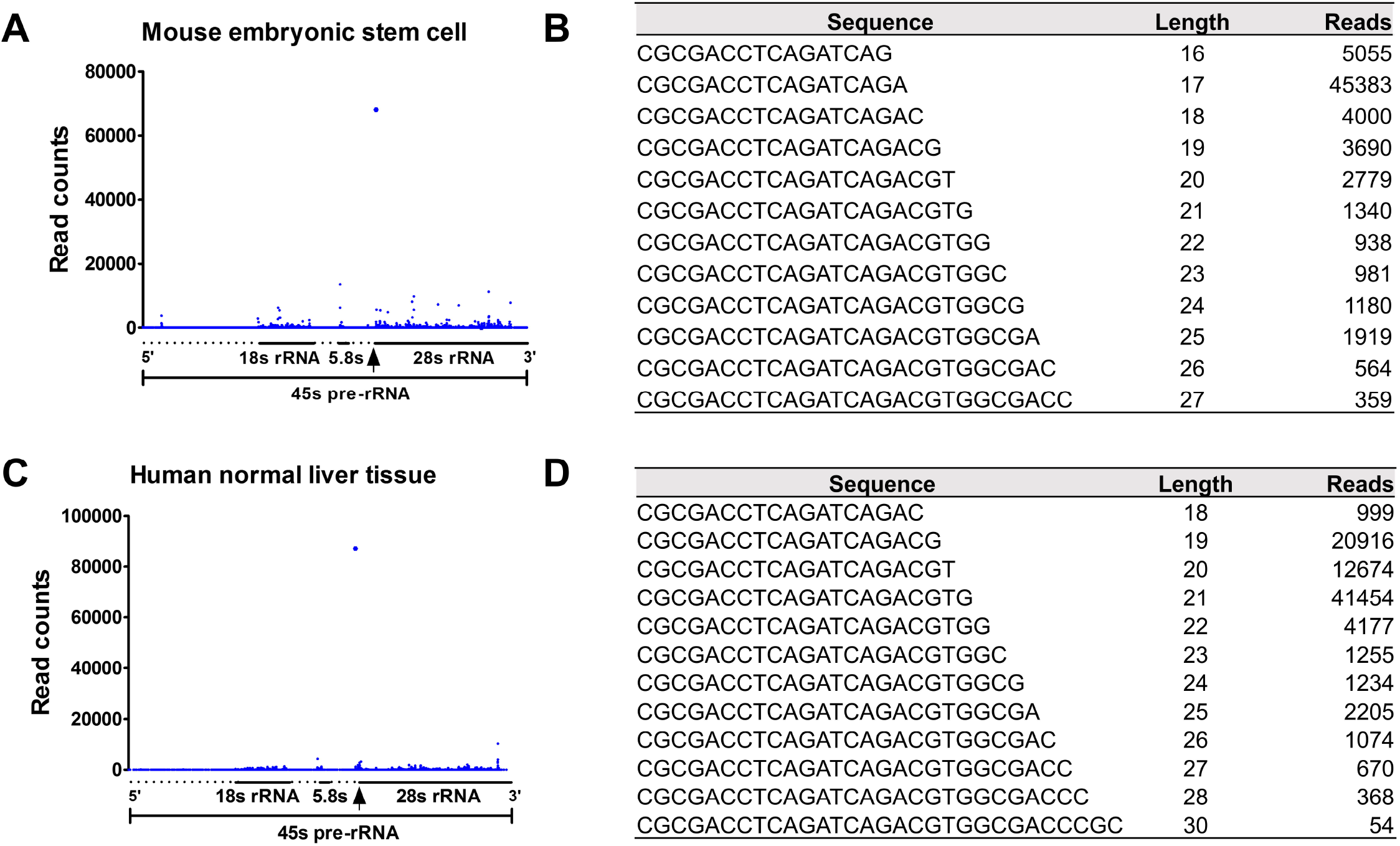
Alignments and statistics of rRNA-derived sRNAs in mouse embryonic stem cells (GSM314552) and human normal liver tissue (GSM531974). sRNA reads from GSM314552 (A) and GSM531974 (C) were aligned along with 45s pre-rRNA. The 5’ end of 28s rRNA was pointed by arrow. Raw read counts were present. Read counts of 28s5-rtsRNAs with different length were showed in (BD).

### 28s5-rtsRNA tissue specificity

To determine tissue specificity of 28s5-rtsRNA, we explore public available human small RNA database DASHR (<u>D</u>atabase of <u>s</u>mall <u>h</u>uman noncoding <u>R</u>NAs), which integrates 187 sRNA deep sequencing datasets from various human tissues (24). To enable comparison across tissues, we employ ‘reads per million’ (RPM), which take into account the library size information for each dataset and is the most commonly used normalization method across different experiments (25). As showed by Figure 2, 28s5-rtsRNA is highly expressed in bladder, monocyte macrophage and skin tissues compared with other tissues.

**Figure 2.**
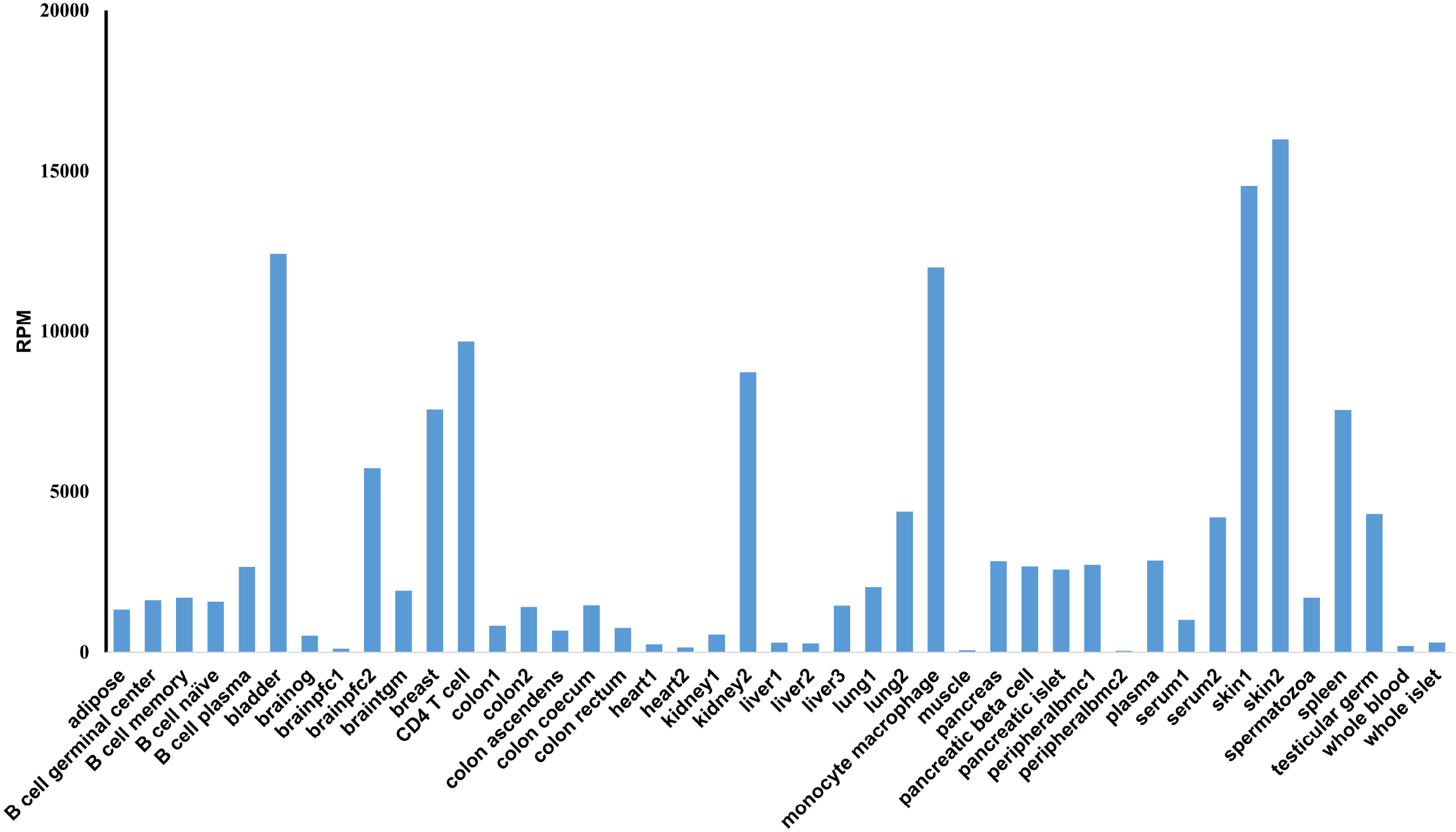
28s5-rtsRNA expression level in different human tissues. sRNA sequencing data from different human tissues was collected, processed, mapped and quantified by DASHR. Read per million (RPM) was present.

Recently, several research showed that tRNA and rRNA served as major resources of sRNA fragments in sperm (13,14,19,26). In mouse, sperm transfer RNA-derived small RNAs (tsRNAs), also known as tRNA fragments (tRFs), mediate intergenerational inheritance of diet-induced metabolic disorders (13,14). These tsRNAs mainly origin from 5’ tRNA with size ranging from 30 to 34 nucleotides. Chu *et al* observed 28s5-rtsRNA accounted for over 1% of total mouse sperm sRNA reads (19). Here, we re-analyze GEO human mature sperm sRNA high-throughput sequencing datasets by using sRNAbench tool, which could conduct small RNA expression profiling, genome mapping and read length statistics (21). Ten sRNA sequencing datasets from four studies were subjected to analysis (Table 1). rRNA-derived small RNA accounts for 2.13% to 13.54% of total assigned small RNA reads in different datasets (Figure 3A). However, tRNA-derived small RNA showed a dramatic dynamics in different studies (form 4.34% to 49.05%, Table 1). This might come from small RNA size fraction during library construction or from sequencing data processing, these lead to the loss of major peak of tsRNA ranging from 30nt to 34nt (Figure 3B). Moreover, rtsRNAs also show a peak around 31nt (SRR8543902, Figure 3B).

**Table 1.** Sperm Small RNA Sequencing Data

| Accession | Study ID | miRNA (%) | rRNA (%) | tRNA (%) | 28s5-rtRNA (RPM) | tRF-Glu-CTC (RPM) | Read Length In Analysis |
| --- | --- | --- | --- | --- | --- | --- | --- |
| SRR1264562 | GSE49624 | 15.05% | 2.13% | 11.79% | 7189 | 41 | 15-26 |
| SRR1264564 | GSE49624 | 8.33% | 9.31% | 9.35% | 57265 | 102 | 15-26 |
| SRR2845753 | GSE74426 | 1.45% | 13.54% | 4.58% | 22735 | 2957 | 15-32 |
| SRR2845754 | GSE74426 | 1.97% | 11.93% | 6.44% | 22910 | 5120 | 15-32 |
| SRR8543902 | GSE110190 | 10.69% | 4.43% | 45.13% | 14235 | 8321 | 15-40 |
| SRR8543903 | GSE110190 | 9.34% | 3.34% | 41.65% | 13531 | 8033 | 15-40 |
| SRR8543925 | GSE110190 | 7.49% | 4.88% | 49.05% | 15373 | 7114 | 15-40 |
| SRR8543926 | GSE110190 | 8.26% | 5.61% | 34.29% | 19083 | 6152 | 15-40 |
| SRR6502268 | GSE109475 | 0.66% | 11.72% | 6.12% | 20317 | 3810 | 15-41 |
| SRR6502271 | GSE109475 | 0.36% | 9.38% | 4.34% | 8914 | 1410 | 15-41 |

**Figure 3.**
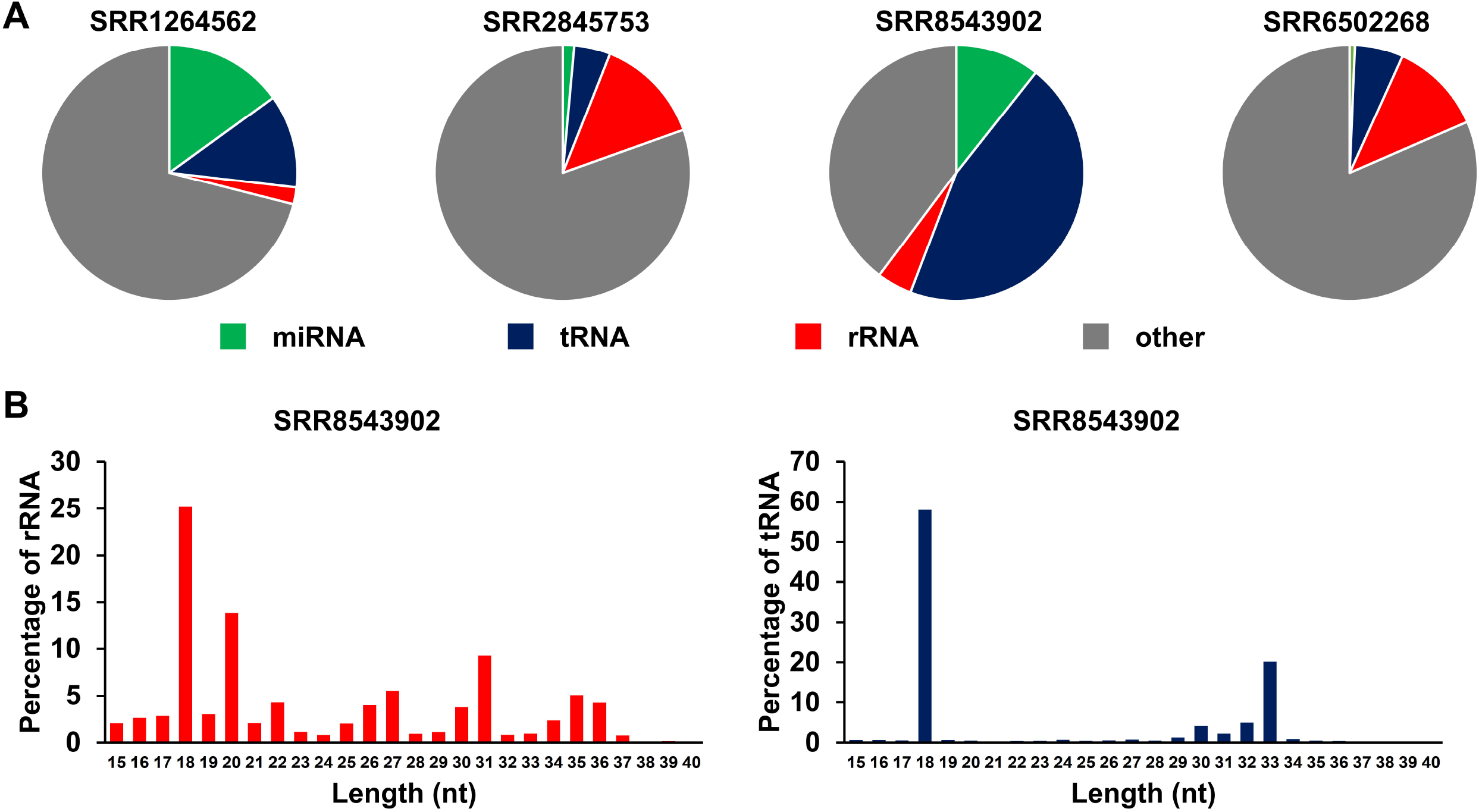
small RNA in sperm. (A) Catalogue of small RNA population in mature human sperm. (B) Length distributions of rRNA and tRNA-derived sRNA in human sperm (SRR8543902).

### 28s5-rtsRNA is independent of canonical miRNA biogenesis pathway

During miRNA maturation, primary miRNA transcripts are successively cleaved by two RNase III Drosha and Dicer. Knockout of Dgcr8 (Drosha partner protein) or Dicer could eliminate miRNA expression in mouse ESCs (23). We reanalyzed these ESCs sRNA sequencing data and found that knockout of Dgcr8 or Dicer have no impact on 28s5-rtsRNA expression (Figure 4AB). Canonical miRNAs, such as let-7c, miR-16 and miR-295, showed a dramatic downregulation after Dgcr8 or Dicer knockout (Figure 4AB). Similarly, knockout of Drosha or Dicer in human HCT116 cell line could not block 28s5-rtsRNA expression (Figure 4CD) (27).

**Figure 4.**
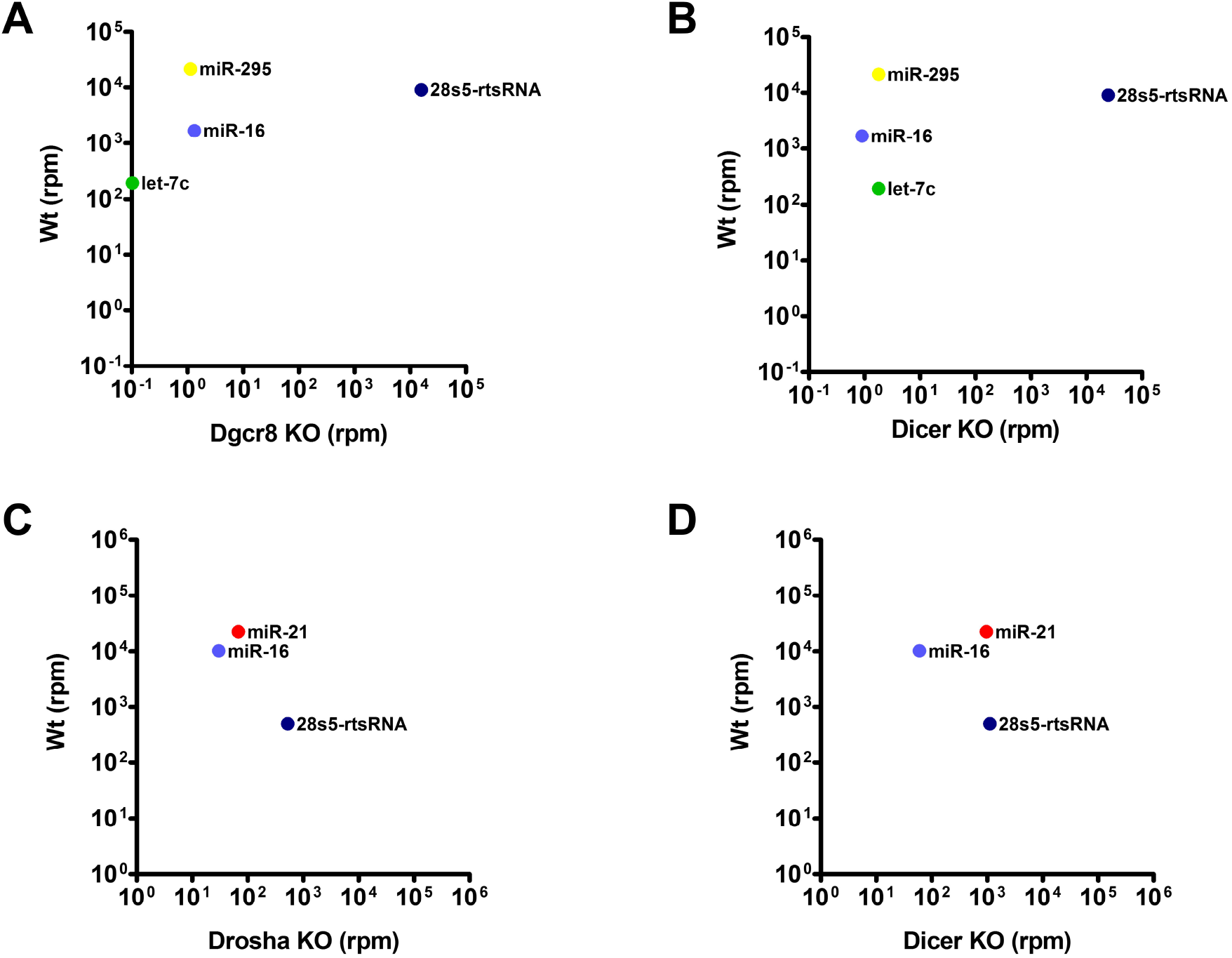
28s5-rtsRNA is independent of miRNA biogenesis pathway. Mouse embryonic stem cells sRNA expression (RPM) in wild-type versus in Dgcr8 KO (A) or in Dicer KO (B) cells. Human HCT116 cell sRNA expression (RPM) in wild-type versus in Drosha KO (C) or in Dicer KO (D) cells.

### 28s5-rtsRNA is not associated with Argonaute protein

In mammalian cells, siRNA and miRNA incorporates into RNA-induced silencing complex (RISC) and programs it to target RNA transcripts (5,28). In RISC, Argonaute (Ago) proteins bind to small RNA and position it in a conformation that facilitates target recognition. Here, we obtain two independent human cell line Ago-immunoprecipitation sRNA deep sequencing data and investigate whether or not 28s5-rtsRNA could incorporate into RISC (29,30). After quality filtering, adapter trimming and length filtering, sRNA expression levels are first present as RPM and then transformed to relative expression (Input is set as one). miRNAs, such as hsa-miR-21-5p, hsa-miR-30a-5p and hsa-miR-92a-3p, are associated with Argonaute proteins, particularly with Ago2 (Figure 5). While for 28s5-rtsRNA, the relative RPM values are very low in Ago-protein groups (Figure 5). This maybe because that Argonaut proteins have a strong preference for uridine at the 5’ end of small RNA.

**Figure 5.**
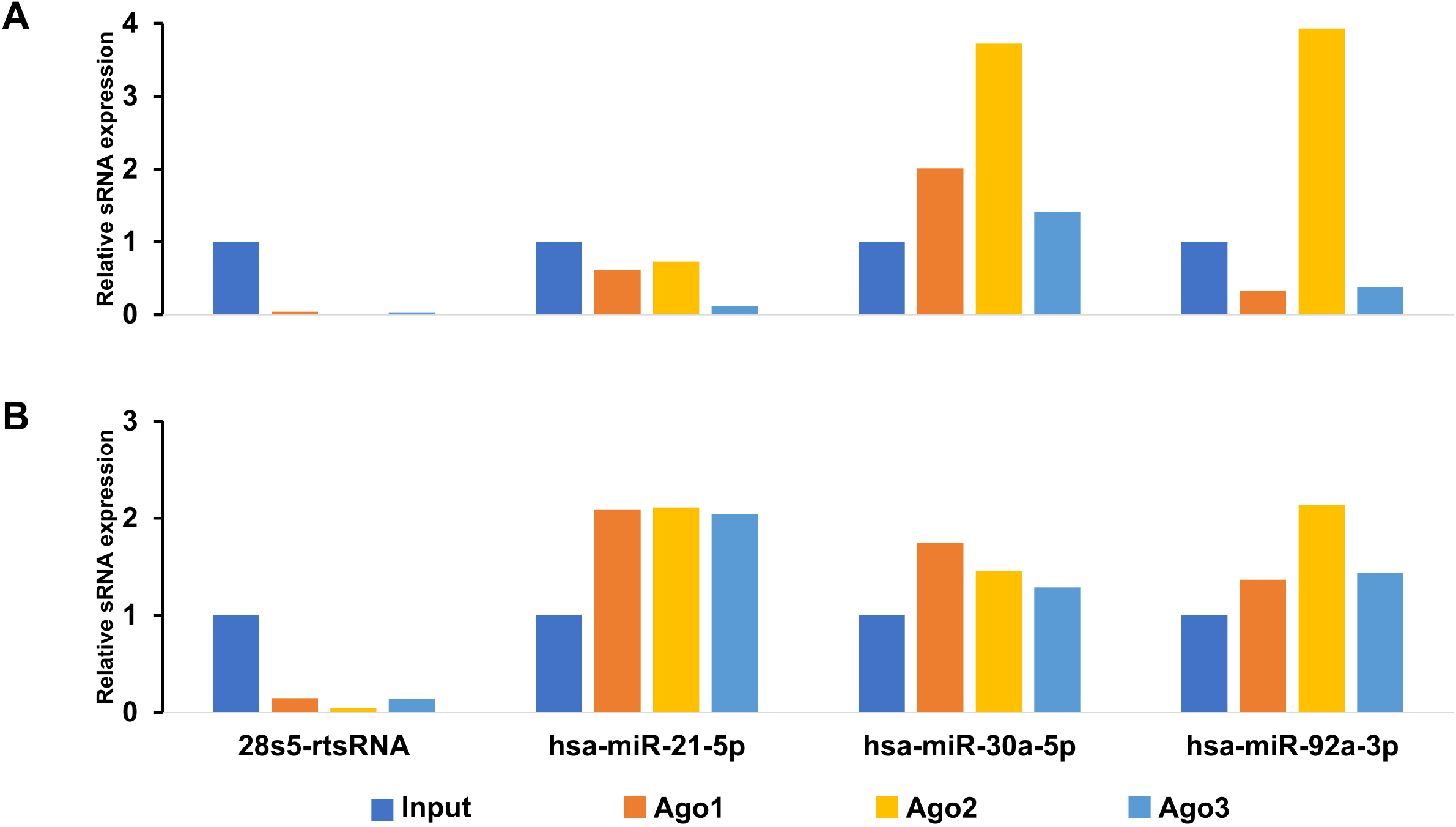
28s5-rtsRNA is not associated with human Argonaut proteins. Two independent Ago-immuneprecipitation sRNA-sequencing data are subject to analysis (A ref. 29) (B ref. 30). Relative 28s5-rtsRNA, has-miR-21-5p, has-miR-30a-5p and has-miR-92a-3p expression levels are present (Inputs set as one).

### Overexpression of 28s5-rtsRNA could reduce the mRNA levels of multiple ribosomal protein

To explore the function of 28s5-rtsRNA, we introduce 28s5-rtsRNA into HeLa cells and analyze the transcriptome dynamics by RNA-sequencing. Negative Control or 28s5-rtsRNA are transfected into HeLa cells. 48 hour after transfection, total RNAs were extracted and subjected to Agilent 2100 sample quality analysis (Supplementary Figure 1A). 28s5-rtsRNA leads to a significant upregulation of 28s/18s rRNA ratio compared with Negative Control transfection group (Supplementary Figure 1B). In the first batch of samples, each group contains two replicates (2 versus 2). We employed DESeq2 method to identify significant differentially expressed genes by setting 1.0E-5 as P-value cutoff (after Benjamini and Hochberg correction) (Supplementary Table 1). Among 189 significant genes, 127 genes are downregulated and 62 genes are upregulated in 28s5-rtsRNA transfection group compared with Negative Control group, respectively. All these 189 genes are also identified as significant differentially expressed genes when using edgeR method setting 1.0E-5 as P-value cutoff (after Benjamini and Hochberg correction) (Supplementary Table 1). We find 11 ribosomal protein large (RPL) and 11 ribosomal protein small (RPS) genes are within downregulated gene list (Supplementary Table 1). We then employ ToppGene Suit (https://toppgene.cchmc.org/) to perform Gene Ontology (GO) analysis. Indeed, GO analysis revealed that the 28s5-rtsRNA-downregulated genes were over represented in gene categories involved in cellular component associated with the ribosome (Table 2).

**Table 2.** Gene Ontology Analysis of 28s5-rtsRNA Downregulated Genes: Cellular Component

| No | ID | Name | pValue | FDR B&H | FDR B&Y | Bonferroni | Genes from Input | Genes in Annotation |
| --- | --- | --- | --- | --- | --- | --- | --- | --- |
| 1 | GO:0022626 | cytosolic ribosome | 2.82E-26 | 1.01E-23 | 6.51E-23 | 1.01E-23 | 22 | 119 |
| 2 | GO:0044391 | ribosomal subunit | 2.05E-22 | 3.66E-20 | 2.37E-19 | 7.32E-20 | 22 | 175 |
| 3 | GO:0005840 | ribosome | 1.42E-19 | 1.27E-17 | 8.20E-17 | 5.08E-17 | 22 | 235 |
| 4 | GO:0044445 | cytosolic part | 1.42E-19 | 1.27E-17 | 8.20E-17 | 5.08E-17 | 22 | 235 |
| 5 | GO:0022627 | cytosolic small ribosomal subunit | 3.28E-15 | 2.01E-13 | 1.30E-12 | 1.17E-12 | 11 | 44 |
| 6 | GO:1990904 | ribonucleoprotein complex | 3.36E-15 | 2.01E-13 | 1.30E-12 | 1.20E-12 | 29 | 745 |
| 7 | GO:0022625 | cytosolic large ribosomal subunit | 4.85E-13 | 2.48E-11 | 1.60E-10 | 1.74E-10 | 11 | 67 |
| 8 | GO:0015935 | small ribosomal subunit | 1.11E-12 | 4.97E-11 | 3.21E-10 | 3.98E-10 | 11 | 72 |
| 9 | GO:0015934 | large ribosomal subunit | 7.66E-11 | 3.05E-09 | 1.97E-08 | 2.74E-08 | 11 | 105 |
| 10 | GO:0005925 | focal adhesion | 6.33E-10 | 2.26E-08 | 1.46E-07 | 2.26E-07 | 17 | 393 |

In the second batch of samples, each group contains six replicates (6 versus 6). DESeq2 statistical test was employed to identify differentially expressed genes by setting 1.0E-20 as P-value cutoff (after Benjamini and Hochberg correction) (Supplementary Table 2). A total of 1103 genes showed a significance between Negative Control group and 28s5-rtsRNA group. We find 26 RPL and 17 RPS genes in 606 28s5-rtsRNA-downregulated genes. GO analysis also showed an over representation of rtsRNA-downregulated genes in ribosome gene categories (Supplementary Table 3). Moreover, we further validated RNA-sequencing results by realtime-qPCR (Figure 6).

**Figure 6.**
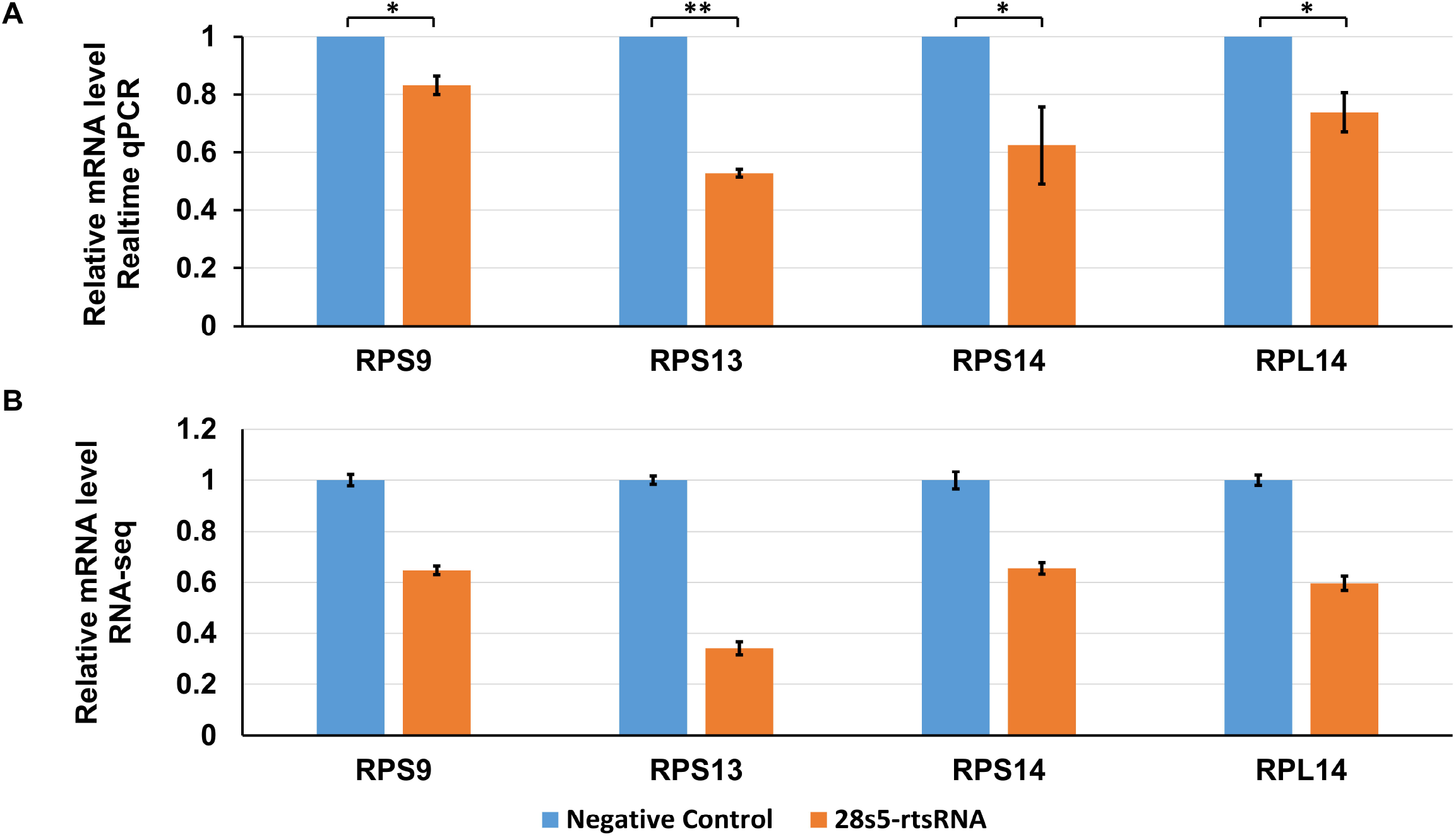
Realtime qPCR validation of RNA-seq results. (A) Realtime qPCR analysis of ribosomal protein mRNA levels in HeLa cells. Relative expression of RPS9, RPS13, RPS14 and RPL14 mRNA were present. GAPDH mRNA levels were used for normalization. (n = 3). (B) Relative gene expression by RNA-sequencing. (n = 6). \**p* < 0.05, \*\**p* < 0.01; statistical significance calculated using two-tailed Student’s t-test.

## Discussion

In present study, we reanalyze the GEO small RNA deep sequencing datasets and find rRNAs serve as major resources of small RNA. In mouse embryotic stem cell and human normal liver tissue, the most abundant rRNA-derived sRNAs are originated from the mature 28s rRNA 5’ ends. Here, we name these 28s rRNA 5’ terminal derived small RNA as 28s5-rtsRNA. By exploring human tissue sRNA sequencing database DASHR, we showed the tissue specificity of 28s5-rtsRNA. We also found 28s5-rtsRNA served as one of the most abundant sRNAs in human sperm. Moreover, we showed 28s5-rtsRNA was independent of miRNA biogenesis pathway and was not associated with Argonaut proteins. At last by RNA-sequencing and realtime-qPCR, we found that 28s5-rtsRNA overexpression could significantly downregulate the expression of multiple ribosomal protein mRNAs. Our results re-emphasized an abundant 5’ 28s rRNA-terminal-derived small RNA (28s5-rtsRNA), which may play a role in synergistic ribosomal protein mRNA expression regulation.

The development of high-throughput sequencing has markedly expanded the categories of sRNA. Recently, ample studies have identified rRNA-derived sRNA, for review see ref (31). We found sRNA from 28s rRNA 5’ terminal end showed a length dynamics with identical 5’ end and different 3’ ends (Figure 1BD). In this study, we assigned these rtsRNA as one type and named them as 28s5-rtsRNA. 28s5-rtsRNA is the most abundant rRNA-derived sRNA (Figure 1AC). Several studies have focused on 28s5-rtsRNA and showed its several special characteristics. Chu *et al* observed a dramatic increase in 28s5-rtsRNA expression in a LPS-induced acute inflammation mouse model (19). By northern blot, Zhang *et al* demonstrated that the knockout of Dnmt2, a multisubstrate tRNA methylatransferase, decreased 28s5-rtsRNA levels in mouse sperm (20). Through analyzing a cDNA library derived from small ribosome-associated RNAs in *Saccharomyces cerevisiae*, Zywicki *et al* detected an abundant 23-nt small RNA, which originated from the 5’-part of 25s rRNA and showed almost exactly identical end (32). However unlike their mammalian counterpart, the 5’ ends of these yeast 25s5-rtsRNA start at the 3rd nucleotide of mature 25s rRNA.

Accumulating evidence from sperm sRNA deep sequencing indicates that housekeeping tRNA and rRNA serve as major resources of sperm sRNA. Here, we reanalyzed multiple human sperm sRNA sequencing datasets from different studies and found 28s5-rtsRNA was among the top abundant sRNAs. While tRNA-derived sRNA accounts for about 40% of total small RNA reads when small RNA fraction size comes to 40nt (Figure 3). The function of sperm 28s5-rtsRNA need to be further investigated. As we showed, 28s5-rtsRNA overexpression could decrease multiple ribosomal protein mRNAs. Thus we hypothesize the abundant sperm 28s5-rtsRNA might help to devoid of ribosome and strip down cytoplasm during spermatogenesis.

Several research has showed that sRNAs could bind to ribosomal protein mRNAs. Kim *et al* found that a specific tRNA-derived sRNA, LeuCAG3’tsRNA, could bind at two ribosomal protein mRNAs (RPS15 and RPS28) to enhance their translation (33). Ørom *et al* employed an affinity-based procedure to obtain the potential targets of miR-10a and found over half of 100 most enriched genes involved in protein biosynthesis and in particular ribosomal proteins (34).

They showed miR-10a could interact with the 5’ untranslated region (5’UTR) of ribosomal protein mRNAs to enhance their translation. Here by RNA-sequencing method, we demonstrate that 28s5-rtsRNA overexpression could significantly decrease multiple ribosomal protein mRNA levels. These results were further validated by realtime-qPCR analysis (Figure 6). However in our RNA-sequencing results, the fold changes of most significant ribosomal protein mRNAs are not less than 0.5 (28s5-rtsRNA overexpression / Negative Control), which is a regular cutoff value of fold change. This maybe because that the ribosomal protein mRNA levels are associated with mass of total RNA. The fold change will be narrowed during RNA-sequencing library construction (equal RNA amount loading) and sequencing data processing process.

In summary, we focus on a novel abundant rRNA-derived small RNA named 28s5-rtsRNA, which is independent of miRNA biogenesis pathway and not associated with Argonaut proteins. Overexpression of 28s5-rtsRNA could decrease multiple ribosomal protein mRNA levels. Our research emphasize 28s5-rtsRNA may act as a negative regulator of ribosomal protein mRNA, thereby mediating the coordinate expression of ribosomal proteins.

## Acknowledgements

This work was supported by the National Natural Science Foundation of China [31870860, 31400673 to S.L.]; the Tianjin Research Program of Application Foundation and Advanced Technology [14JCQNJC09800 to S.L.].

---

**Supplementary Figure 1.**
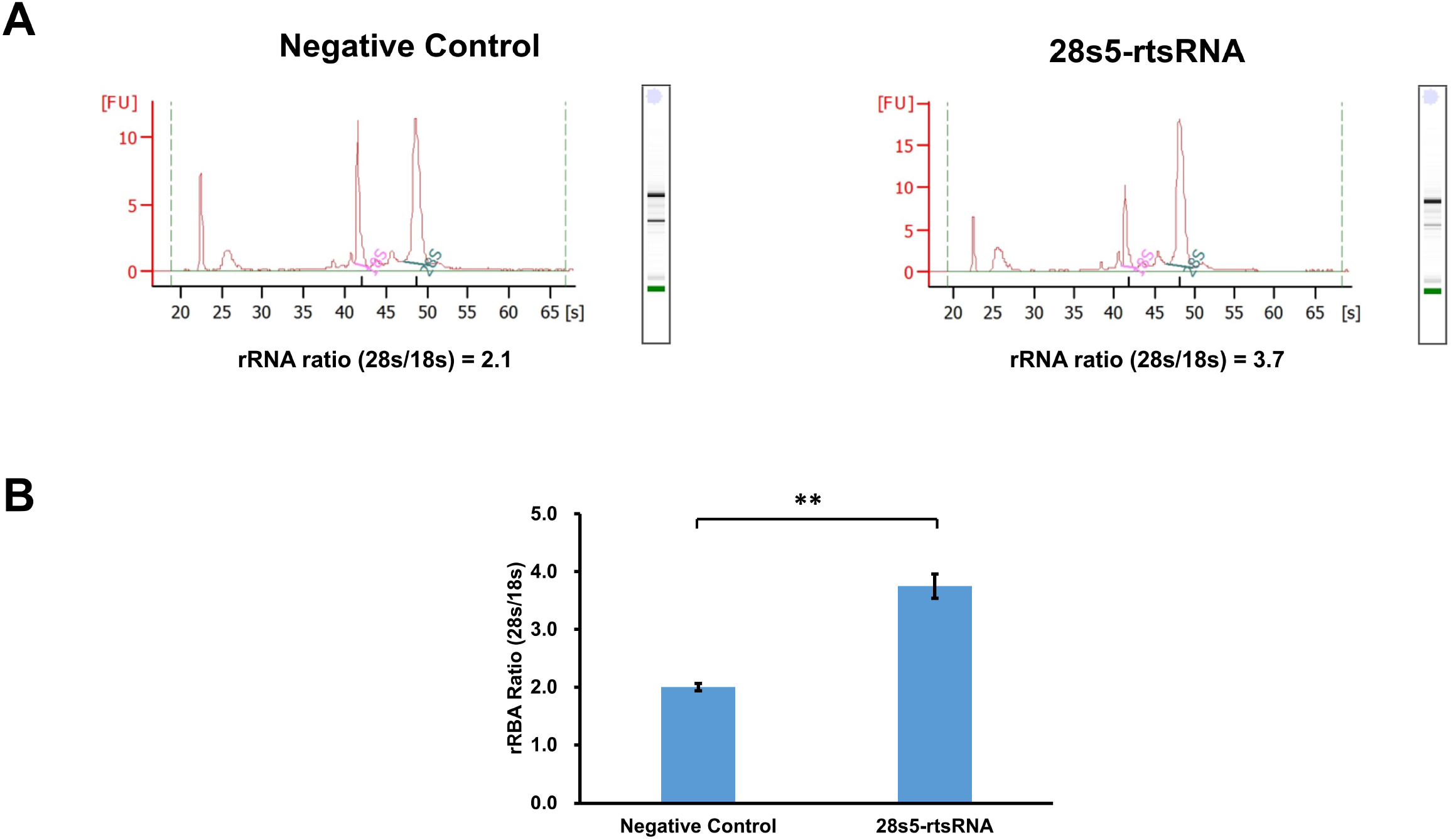
rRNA size distribution analysis. (A) Electrophoretic size distribution of RNAs from Negative Control and 28s5-rtsRNA transfected HeLa cells analyzed by Agilent Bioanalyzer. (B) 28s/18s rRNA ratio in Negative Control and 28s5-trsRNA transfected HeLa cells. (n = 6). \*\**p* < 0.01; statistical significance calculated using two-tailed Student’s t-test.

## References

1. Filipowicz, W., Jaskiewicz, L., Kolb, F.A. and Pillai, R.S. (2005) Post-transcriptional gene silencing by siRNAs and miRNAs. Current opinion in structural biology, 15, 331–341.

2. Moazed, D. (2009) Small RNAs in transcriptional gene silencing and genome defence. Nature, 457, 413–420.

3. Bartel, D.P. (2004) MicroRNAs: genomics, biogenesis, mechanism, and function. Cell, 116, 281–297.

4. Juliano, C., Wang, J. and Lin, H. (2011) Uniting germline and stem cells: the function of Piwi proteins and the piRNA pathway in diverse organisms. Annual review of genetics, 45, 447–469.

5. Liu, Q. and Paroo, Z. (2010) Biochemical principles of small RNA pathways. Annual review of biochemistry, 79, 295–319.

6. Shi, W., Hendrix, D., Levine, M. and Haley, B. (2009) A distinct class of small RNAs arises from pre-miRNA-proximal regions in a simple chordate. Nature structural & molecular biology, 16, 183–189.

7. Ender, C., Krek, A., Friedlander, M.R., Beitzinger, M., Weinmann, L., Chen, W., Pfeffer, S., Rajewsky, N. and Meister, G. (2008) A human snoRNA with microRNA-like functions. Molecular cell, 32, 519–528.

8. Taft, R.J., Glazov, E.A., Lassmann, T., Hayashizaki, Y., Carninci, P. and Mattick, J.S. (2009) Small RNAs derived from snoRNAs. RNA, 15, 1233–1240.

9. Fu, H., Feng, J., Liu, Q., Sun, F., Tie, Y., Zhu, J., Xing, R., Sun, Z. and Zheng, X. (2009) Stress induces tRNA cleavage by angiogenin in mammalian cells. FEBS letters, 583, 437–442.

10. Lee, Y.S., Shibata, Y., Malhotra, A. and Dutta, A. (2009) A novel class of small RNAs: tRNA-derived RNA fragments (tRFs). Genes & development, 23, 2639–2649.

11. Cole, C., Sobala, A., Lu, C., Thatcher, S.R., Bowman, A., Brown, J.W., Green, P.J., Barton, G.J. and Hutvagner, G. (2009) Filtering of deep sequencing data reveals the existence of abundant Dicer-dependent small RNAs derived from tRNAs. RNA, 15, 2147–2160.

12. Li, Z., Ender, C., Meister, G., Moore, P.S., Chang, Y. and John, B. (2012) Extensive terminal and asymmetric processing of small RNAs from rRNAs, snoRNAs, snRNAs, and tRNAs. Nucleic acids research, 40, 6787–6799.

13. Chen, Q., Yan, M., Cao, Z., Li, X., Zhang, Y., Shi, J., Feng, G.H., Peng, H., Zhang, X., Qian, J. et al. (2016) Sperm tsRNAs contribute to intergenerational inheritance of an acquired metabolic disorder. Science, 351, 397–400.

14. Sharma, U., Conine, C.C., Shea, J.M., Boskovic, A., Derr, A.G., Bing, X.Y., Belleannee, C., Kucukural, A., Serra, R.W., Sun, F. et al. (2016) Biogenesis and function of tRNA fragments during sperm maturation and fertilization in mammals. Science, 351, 391–396.

15. Watson, E.D. (2016) Transferring Fragments of Paternal Metabolism to the Offspring. Cell metabolism, 23, 401–402.

16. Shi, J., Ko, E.A., Sanders, K.M., Chen, Q. and Zhou, T. (2018) SPORTS1.0: A Tool for Annotating and Profiling Non-coding RNAs Optimized for rRNA- and tRNA-derived Small RNAs. Genomics, proteomics & bioinformatics, 16, 144–151.

17. Lee, H.C., Chang, S.S., Choudhary, S., Aalto, A.P., Maiti, M., Bamford, D.H. and Liu, Y. (2009) qiRNA is a new type of small interfering RNA induced by DNA damage. Nature, 459, 274–277.

18. Wei, H., Zhou, B., Zhang, F., Tu, Y., Hu, Y., Zhang, B. and Zhai, Q. (2013) Profiling and identification of small rDNA-derived RNAs and their potential biological functions. PloS one, 8, e56842.

19. Chu, C., Yu, L., Wu, B., Ma, L., Gou, L.T., He, M., Guo, Y., Li, Z.T., Gao, W., Shi, H. et al. (2017) A sequence of 28S rRNA-derived small RNAs is enriched in mature sperm and various somatic tissues and possibly associates with inflammation. Journal of molecular cell biology, 9, 256–259.

20. Zhang, Y., Zhang, X., Shi, J., Tuorto, F., Li, X., Liu, Y., Liebers, R., Zhang, L., Qu, Y., Qian, J. et al. (2018) Dnmt2 mediates intergenerational transmission of paternally acquired metabolic disorders through sperm small non-coding RNAs. Nature cell biology, 20, 535–540.

21. Rueda, A., Barturen, G., Lebron, R., Gomez-Martin, C., Alganza, A., Oliver, J.L. and Hackenberg, M. (2015) sRNAtoolbox: an integrated collection of small RNA research tools. Nucleic acids research, 43, W467–473.

22. Chen, J., Bardes, E.E., Aronow, B.J. and Jegga, A.G. (2009) ToppGene Suite for gene list enrichment analysis and candidate gene prioritization. Nucleic acids research, 37, W305–311.

23. Babiarz, J.E., Ruby, J.G., Wang, Y., Bartel, D.P. and Blelloch, R. (2008) Mouse ES cells express endogenous shRNAs, siRNAs, and other Microprocessor-independent, Dicer-dependent small RNAs. Genes & development, 22, 2773–2785.

24. Leung, Y.Y., Kuksa, P.P., Amlie-Wolf, A., Valladares, O., Ungar, L.H., Kannan, S., Gregory, B.D. and Wang, L.S. (2016) DASHR: database of small human noncoding RNAs. Nucleic acids research, 44, D216–222.

25. Anders, S., McCarthy, D.J., Chen, Y., Okoniewski, M., Smyth, G.K., Huber, W. and Robinson, M.D. (2013) Count-based differential expression analysis of RNA sequencing data using R and Bioconductor. Nature protocols, 8, 1765–1786.

26. Peng, H., Shi, J., Zhang, Y., Zhang, H., Liao, S., Li, W., Lei, L., Han, C., Ning, L., Cao, Y. et al. (2012) A novel class of tRNA-derived small RNAs extremely enriched in mature mouse sperm. Cell research, 22, 1609–1612.

27. Kim, Y.K., Kim, B. and Kim, V.N. (2016) Re-evaluation of the roles of DROSHA, Export in 5, and DICER in microRNA biogenesis. Proceedings of the National Academy of Sciences of the United States of America, 113, E1881–1889.

28. Pratt, A.J. and MacRae, I.J. (2009) The RNA-induced silencing complex: a versatile gene-silencing machine. The Journal of biological chemistry, 284, 17897–17901.

29. Dueck, A., Ziegler, C., Eichner, A., Berezikov, E. and Meister, G. (2012) microRNAs associated with the different human Argonaute proteins. Nucleic acids research, 40, 9850–9862.

30. Burroughs, A.M., Ando, Y., de Hoon, M.J., Tomaru, Y., Suzuki, H., Hayashizaki, Y. and Daub, C.O. (2011) Deep-sequencing of human Argonaute-associated small RNAs provides insight into miRNA sorting and reveals Argonaute association with RNA fragments of diverse origin. RNA biology, 8, 158–177.

31. Lambert, M., Benmoussa, A. and Provost, P. (2019) Small Non-Coding RNAs Derived From Eukaryotic Ribosomal RNA. Non-coding RNA, 5.

32. Zywicki, M., Bakowska-Zywicka, K. and Polacek, N. (2012) Revealing stable processing products from ribosome-associated small RNAs by deep-sequencing data analysis. Nucleic acids research, 40, 4013–4024.

33. Kim, H.K., Fuchs, G., Wang, S., Wei, W., Zhang, Y., Park, H., Roy-Chaudhuri, B., Li, P., Xu, J., Chu, K. et al. (2017) A transfer-RNA-derived small RNA regulates ribosome biogenesis. Nature, 552, 57–62.

34. Orom, U.A., Nielsen, F.C. and Lund, A.H. (2008) MicroRNA-10a binds the 5’UTR of ribosomal protein mRNAs and enhances their translation. Molecular cell, 30, 460–471.

