## Supplementary Table 3 for "Human 28s rRNA 5’ terminal derived small RNA inhibits ribosomal protein mRNA levels"

**Supplementary Table 3. GO Analysis of 28s5-rtsRNA Downregulated Genes: Cellular Component**

| No | ID | Name | pValue | FDR B&H | FDR B&Y | Bonferroni | Genes from Input | Genes in Annotation |
| --- | --- | --- | --- | --- | --- | --- | --- | --- |
| 1 | GO:0022626 | cytosolic ribosome | 1.08E-33 | 6.88E-31 | 4.84E-30 | 6.88E-31 | 42 | 119 |
| 2 | GO:0044391 | ribosomal subunit | 3.98E-27 | 1.27E-24 | 8.96E-24 | 2.55E-24 | 43 | 175 |
| 3 | GO:0044445 | cytosolic part | 1.41E-22 | 3.00E-20 | 2.11E-19 | 9.00E-20 | 44 | 235 |
| 4 | GO:0005840 | ribosome | 1.11E-21 | 1.78E-19 | 1.25E-18 | 7.11E-19 | 43 | 235 |
| 5 | GO:0022625 | cytosolic large ribosomal subunit | 3.11E-21 | 3.98E-19 | 2.80E-18 | 1.99E-18 | 25 | 67 |
| 6 | GO:0005925 | focal adhesion | 3.09E-17 | 3.30E-15 | 2.32E-14 | 1.98E-14 | 49 | 393 |
| 7 | GO:0005924 | cell-substrate adherens junction | 5.20E-17 | 4.76E-15 | 3.35E-14 | 3.33E-14 | 49 | 398 |
| 8 | GO:0030055 | cell-substrate junction | 8.68E-17 | 6.94E-15 | 4.89E-14 | 5.56E-14 | 49 | 403 |
| 9 | GO:0015934 | large ribosomal subunit | 6.19E-16 | 4.41E-14 | 3.10E-13 | 3.96E-13 | 25 | 105 |
| 10 | GO:0022627 | cytosolic small ribosomal subunit | 3.73E-15 | 2.39E-13 | 1.68E-12 | 2.39E-12 | 17 | 44 |
